# Bacterial Microtubules Exhibit Polarized Growth, Mixed-Polarity Bundling, and Destabilization by GTP Hydrolysis

**DOI:** 10.1101/112987

**Authors:** César Díaz-Celis, Viviana I. Risca, Felipe Hurtado, Jessica K. Polka, Scott D. Hansen, Daniel Maturana, Rosalba Lagos, R. Dyche Mullins, Octavio Monasterio

## Abstract

Bacteria of the genus *Prosthecobacter* express homologs of eukaryotic α-and β-tubulin, called BtubA and BtubB, that have been observed to assemble into bacterial microtubules (bMTs). The *btubAB* genes likely entered the *Prosthecobacter* lineage via horizontal gene transfer and may derive from an early ancestor of the modern eukaryotic microtubule (MT). Previous biochemical studies revealed that BtubA/B polymerization is GTP-dependent and reversible and that BtubA/B folding does not require chaperones. To better understand bMT behavior and gain insight into the evolution of microtubule dynamics, we characterized *in vitro* bMT assembly using a combination of polymerization kinetics assays, and microscopy. Like eukaryotic microtubules, bMTs exhibit polarized growth with different assembly rates at each end. GTP hydrolysis stimulated by bMT polymerization drives a stochastic mechanism of bMT disassembly that occurs via polymer breakage. We also observed treadmilling (continuous addition and loss of subunits at opposite ends) of bMT fragments. Unlike MTs, polymerization of bMTs requires KCl, which reduces the critical concentration for BtubA/B assembly and induces bMTs to form stable mixed-orientation bundles in the absence of any additional bMT-binding proteins. Our results suggest that at potassium concentrations resembling that inside the cytoplasm of *Prosthecobacter*, bMT stabilization through self-association may be a default behavior. The complex dynamics we observe in both stabilized and unstabilized bMTs may reflect common properties of an ancestral eukaryotic tubulin polymer.

**Importance:** Microtubules are polymers within all eukaryotic cells that perform critical functions: they segregate chromosomes in cell division, organize intracellular transport by serving as tracks for molecular motors, and support the flagella that allow sperm to swim. These functions rely on microtubules remarkable range of tunable dynamic behaviors. Recently discovered bacterial microtubules composed of an evolutionarily related protein are evolved from a missing link in microtubule evolution, the ancestral eukaryotic tubulin polymer. Using microscopy and biochemical approaches to characterize bacterial microtubules, we observed that they exhibit complex and structurally polarized dynamic behavior like eukaryotic microtubules, but differ in how they self-associate into bundles and become destabilized. Our results demonstrate the diversity of mechanisms that microtubule-like filaments employ to promote filament dynamics and monomer turnover.

## Introduction

BtubA/B is the first prokaryotic tubulin homolog that has been reported to form a microtubule-like structure, called the bacterial microtubule (bMT) in bacterial cells (1). bMTs are assembled from BtubA/B heterodimers (2–4), made of bacterial tubulin A and bacterial tubulin B (BtubA and BtubB), the closest known prokaryotic homologs of eukaryotic tubulin (~35% sequence identity with α/β-tubulin) (1). *btubA* and *btubB* genes were discovered, along with a kinesin light chain homolog (*bklc*) in the genome of four *Prosthecobacter* bacterial species (5), to form the bacterial tubulin operon (btub-operon) (6). Because of this high sequence identity and structural similarity, it has been proposed that *Prosthecobacter* acquired the *btub*-operon by horizontal gene transfer (2, 4–9) from an unknown species. Although the function of bMTs in *Prosthecobacter* remains unknown, they may contribute to the elongated shape of the *Prosthecobacter* cell (4), or serve as a scaffold for the formation of the cell stalk (7).

bMTs observed *in vivo* using cryo-electron tomography have been proposed to consist of 7.6 nm-wide hollow tubes formed by 5 protofilaments (1). Single BtubA/B filaments formed *in vitro* and observed using negatively stained electron microscopy (EM) have been interpreted of consisting of two parallel protofilaments, although some filament bundles have appeared to form a lumen (4). Based on results of pelleting assays (2, 4), and mutagenesis studies (3), it has been inferred that bMT protofilaments contain a polar, alternating linear arrangement of BtubA/B heterodimers. By comparison, eukaryotic MTs are tubular, 25 nm wide polymers formed by about 14 protofilaments, each of which is a polar chain of α/β-tubulin heterodimers (10). MTs are also kinetically polar, with different assembly/disassembly rates at the two ends (11, 12). Polymerization studies have not yet established whether the two bMT ends polymerize at different rates.

All tubulin family proteins, including eukaryotic tubulins, the better characterized prokaryotic tubulins FtsZ and TubZ, and the newly discovered bacteriophage tubulin PhuZ, are assembly-dependent GTPases that polymerize when GTP-bound and are destabilized by hydrolysis of GTP to GDP (13, 14). Similar to α/β-tubulin and FtsZ assembly, BtubA/B polymerization is GTP-dependent, cooperative and reversible, and GTP hydrolysis is coupled to assembly (2, 4, 7). Structural changes in the polymer associated with GTP hydrolysis give rise to complex behaviors in the tubulin family, including dynamic instability observed in eukaryotic MTs (stochastic transitions between growing and shortening phases at microtubule ends (10, 11, 15)), and treadmilling observed in several types of tubulin family polymers (concomitant assembly at one end and disassembly at the other (10)). Whether BtubA/B filaments exhibit any of these or other dynamic behaviors remains an open question.

In order to obtain insight into the dynamics and polarity of BtubA/B polymers, we characterized their assembly *in vitro* with confocal fluorescence microscopy, electron microscopy, light scattering, high-speed pelleting and GTPase assays. We determined that, like eukaryotic MTs, *in vitro*-polymerzed BtubA/B filaments are kinetically polar polymers that are destabilized by GTP hydrolysis. They exhibit stochastic growth and disassembly and treadmilling. Salt stimulates BtubA/B polymerization by reducing the critical concentration for assembly. Unlike eukaryotic MTs, bMTs form stable apolar bundles in the presence of high potassium concentration without any bundling or stabilizing microtubule-binding proteins.

## Results

### Potassium alters the critical concentration of BtubA/B assembly and induces bundling

To date, two methods of purifying recombinant BtubA/B from the *E. coli* soluble cell fraction have been described. The first is high-level expression of the *btubAB* bicistronic unit under the control of T7 promoter, followed by protein purification with anion-exchange and size exclusion chromatography (2). Recently, an additional polymerization and depolymerization cycle step was added to this procedure (7, 16). The second method is the expression of N-terminal(3, 4) or C-terminal (1) (His)_6_-tagged BtubA and BtubB followed by Ni-NTA affinity chromatography, making it possible to purify BtubA and BtubB separately, as well as mutants with impaired assembly. However, (His)_6_-tags can induce the self-assembly of BtubB rings (3, 4). In order to avoid the use of (His)_6_-tagged proteins and simplify the purification process, we designed a new purification method for untagged recombinant BtubA/B, based on the standard α/β-tubulin method of successive cycles of polymerization-depolymerization (see Fig. S1 in the supplemental material). BtubA/B polymerization was in the presence of MgCl_2_ and KCl, which are known to be necessary for polymerization (2, 7). Interestingly, it was not possible to purify BtubA/B using NaCl instead of KCl, and increasing the KCl concentration increased the yield. This suggested to us that potassium regulates a yet uncharacterized BtubA/B assembly parameter.

To determine the effect of KCl on BtubA/B polymer assembly, we used confocal microscopy to visualize GTP-induced polymerization of fluorescently labeled BtubA/B (Fig. 1A). In the absence of KCl, we did not observe discernible filamentous structures. Short and dispersed BtubA/B filaments are visible at 50 mM KCl, and a network of BtubA/B filament bundles appear at 100 mM KCl. At 1 M KCl, bundles increase in number but decrease in length, preventing the formation of a network of bundles. The observed decrease in bundle length is likely due to the large number of bMT ends, which quickly use up the protein available for bMT elongation.

**Fig. 1.**
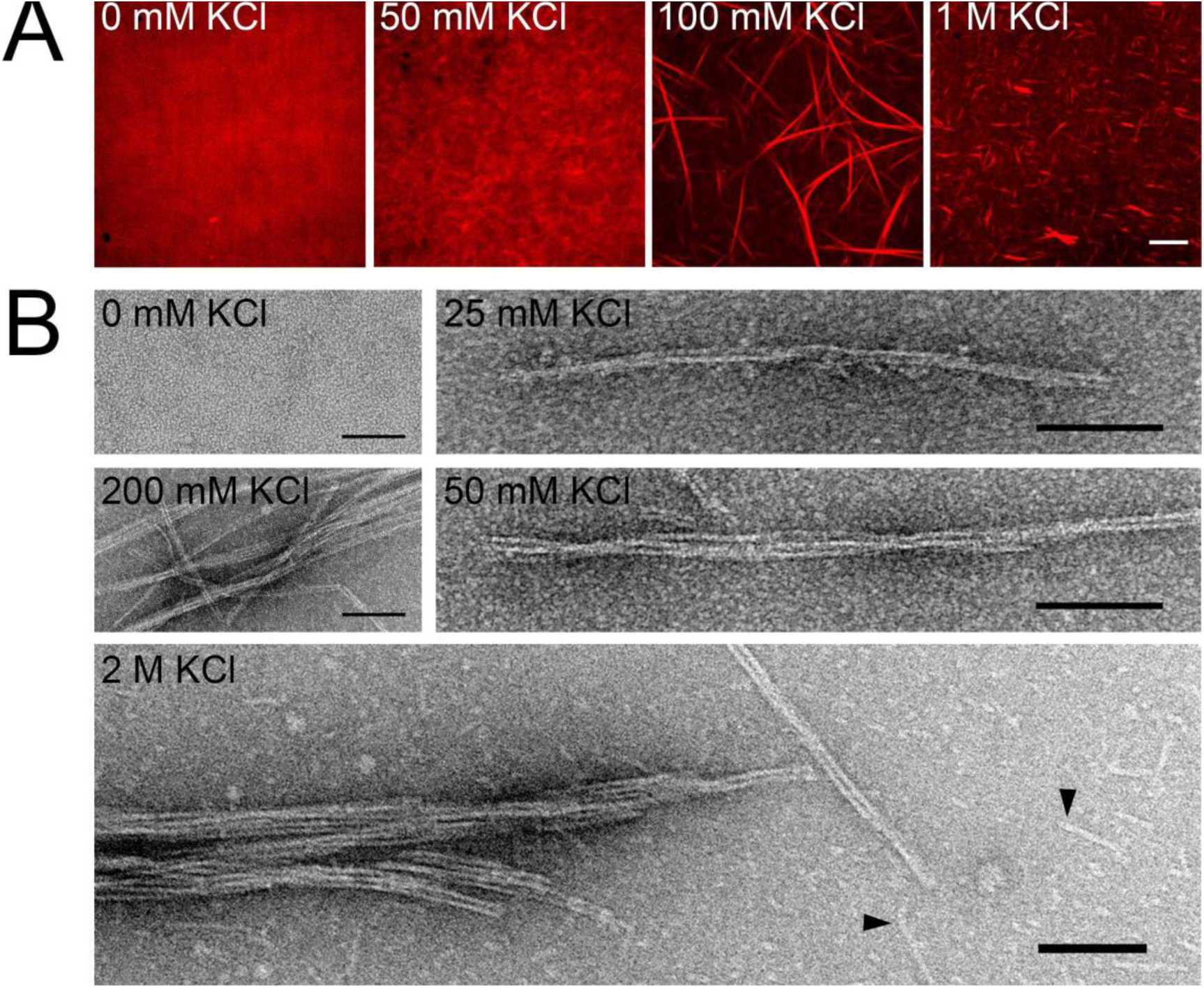
BtubA/B filament morphology and KCl-dependent bundling. A) 5 μM BtubA/B (15% TAMRA-labeled) was polymerized with 1 mM GTP in polymerization buffer (50 mM HEPES-KOH (pH 7), 5 mM MgCl2) with varying KCl concentrations. After 5 min, the samples were applied to a poly-L-lysine coated coverslip and visualized by confocal microscopy. (B) 5 μM BtubA/B was polymerized with 1 mM GTP for 5 min at room temperature in varying KCl concentrations. BtubA/B filaments (bMTs) were stained with uranyl formate 0.75% for EM. bMTs were not observed at 0 M KCl. Above 50 mM KCl, bMTs associated to form bundles. At 2 M KCl, bundles coexist with single bMTs and small structures (black arrowheads). (Scale bar: 10 μm in (A); 100 nm in (B).)

Using negative stain EM, we observed in 25 mM KCl what appeared to be a single filament composed of a pair of BtubA/B protofilaments with a narrow dark space between them (Fig. 1B). We will refer to these filaments as bMTs. Their appearance is similar to paired protofilaments previously observed *in vitro* (4), but may also be consistent with a projection of the five-protofilament bMTs observed by cryo-electron microscopy *in vivo* (1), because the width of the dark space between the filaments is similar to the width of the lumen. Bundling observable by EM at different salt concentrations was consistent with fluorescence microscopy results (Fig. 1A). We observe multiple paired protofilaments in bundles, but not three-protofilament bMTs as previously seen *in vitro* (4). At 2M KCl, we also observed short linear structures, smaller than a single bMT, which were absent below 50 mM KCl (Fig. 1B).

We characterized the effect of potassium on BtubA/B assembly kinetics by 90° light scattering (Fig. 2). Confocal and electron microscopy shows that the increase of KCl concentration induces the bundling of bMTs. Because light scattering depends on the size and shape of the polymers, a deviation of the sigmoidal kinetic behavior reflects a high-order filament association such as bundling (17). bMT assembly kinetics in HEPES buffer with trace KOH showed a lag phase followed by elongation, a peak of the intensity signal, and a decrease due to depolymerization (Fig. 2A). The length of the lag phase is inversely proportional to BtubA/B concentration, confirming that bMT assembly is a nucleation-elongation process, as has been previously suggested (4). At 3.1, 6.3, (see Fig. S2A and S2B in the supplemental material) and 12.5 mM KCl (Fig. 2B), assembly traces were sigmoidal and the lowest BtubA/B concentration at which polymerization was detected decreased with increasing salt. In the range of 25-50 mM KCl, the assembly traces were not sigmoidal and depolymerization was slower (see Fig. S2C and S2D in the supplemental material). This non-sigmoidal assembly behavior may be related to the onset of filament bundling. At 100 (Fig. 2C) and 200 mM KCL (see Fig. S2E in the supplemental material), BtubA/B assembly was fast, and the elongation phase appeared to have a fast step followed by a second slower step that reached the maximum intensity signal or steady state. At 500 mM KCl (Fig. 2D), the light scattering signal was stable for more than 50 min, in contrast to rapidly-decaying signal from BtubA/B polymerized under 25 mM KCl. At ≥1 M KCl, BtubA/B assembly showed an overshoot (see Fig. S2F and S2G in the supplemental material).

**Fig. 2.**
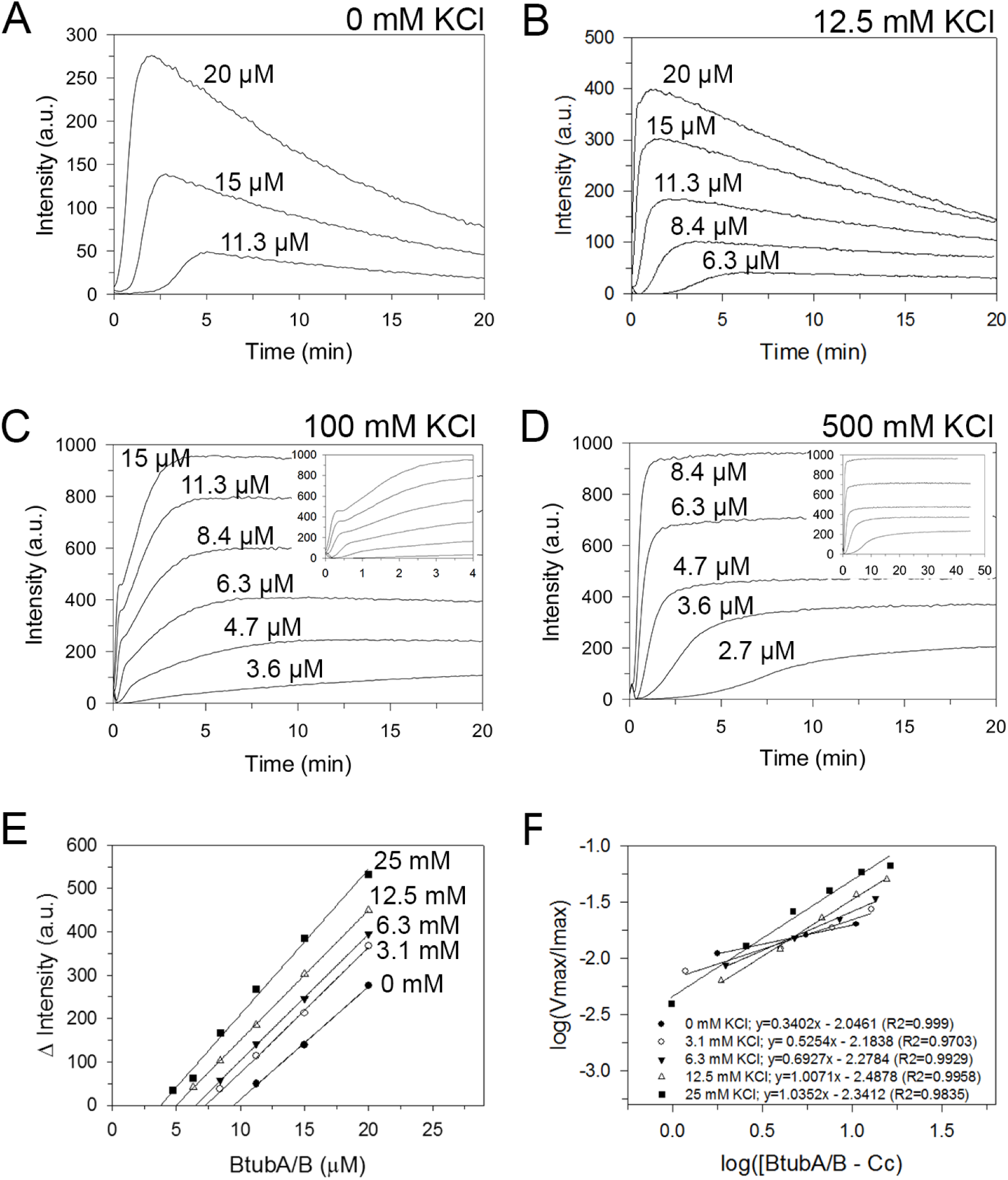
Light scattering of BtubA/B polymerization. Potassium affects bMT assembly kinetics and reduces the critical concentration for BtubA/B assembly. (A) Time course of BtubA/B polymerization monitored by light scattering. Serially diluted BtubA/B (concentrations indicated in the figure) was polymerized by adding 1 mM GTP at 25 °C,in absence of KCl in the polymerization buffer (50 mM HEPES (pH 7), 5 mM MgCl_2_), and with (B) 12.5 mM KCl, (C) 100 mM KCl, and (D) 500 mM KCl. Inset of (C): expanded view of early polymerization. Inset of (D): longer time course. (E) Estimation of the critical concentration of BtubA/B polymerization below 25 mM KCl. The maximum light scattering intensity from each polymerization time course in each condition is plotted versus the initial BtubA/B concentration. The x-intercept of each linear fit corresponds to the critical concentration. (F) Determination of the bMT nucleus size between 0 and 25 mM KCl shows that the rise of KCl concentration increased the size of the polymerization nucleus. The log of the maximum rate of assembly (V_max_) at each KCl condition was normalized by the maximum intensity (I_max_) and was plotted *versus* the log of the BtubA/B available to polymerize (total BtubA/B minus critical concentration). The slope of the linear fit correspond to the half of the nucleus size (*n*/2) (20). Inset: linear fit equation where “x” correspond to the slope and “R2” correspond to the R-squared.

We observed that the protein concentration at which bMT assembly was first detected decrease with increasing KCl, a trend observed over all KCl concentrations employed. This suggests that KCl affects not only bMT bundling, but also bMT formation. Because light scattering is proportional to the amount of assembled polymer and its spatial organization (18)), we determined the critical concentration of BtubA/B assembly from the plot of the maximum scattering intensity *versus* BtubA/B concentration (Fig. 2E) (12, 19). However, we only used data between 0 and 25 mM KCL to avoid the confounding effects of bundling. From the x-axis intersections of Fig. 2E, we determined that in the absence of KCl, the critical concentration for BtubA/B assembly of bMTs is 9.5 μM, and it decreases to 3.8 μM at 25 mM KCl (Table 1). To determine if KCl has an effect on BtubA/B nucleation in this low salt regime, we normalized the log of the maximum rate of assembly (V_max_) by the maximum intensity (I_max_) of each polymerization trace, and this value was plotted *versus* the log of the BtubA/B available to polymerize (Total BtubA/B minus critical concentration of BtubA/B assembly) (Fig. 2F). The slope of the linear fit is equivalent to the half of the nucleus size (*n*/2) (20) and suggests that the rise of KCl concentration increased the size of the BtubA/B polymerization nucleus. At 0 mM KCl, the slope of the linear fit is 0.3, suggesting that the nucleus consist of just a monomer (BtubA or BtubB). At 3.1 mM KCl the nucleus is composed by one BtubA/B heterodimer (*n*=~0.5), and at 12.5-25 mM KCl the slope of the linear fit (*n*=~1) suggest that the nucleus is composed of two BtubA/B heterodimers.

**Table 1.** Critical concentration of BtubA/B assembly, and GTPase activity in different potassium concentrations. Critical concentration of BtubA/B polymerization determined by light scattering and high-speed pelleting, and BtubA/B GTPase activity at different KCl concentrations. ND, not determined.

| KCl (mM) | Steady state [BtubA/B]<br>(Light scattering, $\mu\text{M}$)<br>n=1 | Steady state<br>[BtubA/B] (High-speed pelleting,<br>$\mu\text{M}$)<br>n=3 | GTPase activity (mol<br>GTP $\text{min}^{-1}$ mol<br>BtubA/B$^{-1}$)<br>n=3 |
| --- | --- | --- | --- |
| 0 | 9.5 | $7.2 \pm 1.8$ | $0.96 \pm 0.03$ |
| 3.1 | 7.2 | ND | $1.09 \pm 0.04$ |
| 6.3 | 6.5 | $4.7 \pm 2.4$ | $1.17 \pm 0.08$ |
| 12.5 | 5.0 | $2.4 \pm 1.3$ | $1.29 \pm 0.03$ |
| 25 | 3.8 | $3.2 \pm 0.9$ | $1.36 \pm 0.11$ |
| 50 | ND | $2.9 \pm 0.3$ | $1.32 \pm 0.16$ |
| 100 | ND | $3.0 \pm 0.3$ | $1.0 \pm 0.04$ |
| 200 | ND | $2.6 \pm 0.4$ | $0.95 \pm 0.04$ |
| 500 | ND | $2.2 \pm 0.2$ | $0.58 \pm 0.10$ |
| 1,000 | ND | $1.5 \pm 0.2$ | $0.38 \pm 0.08$ |
| 2,000 | ND | $1.3 \pm 0.3$ | $0.3 \pm 0.02$ |

As an independent measurement, we determined the amount of assembled bMTs and the critical concentration of BtubA/B assembly as a function of KCl concentration using high speed-pelleting. The increment from 0 to 2 M KCl produced an increase of the assembled polymer mass (Fig. 3A) and reduce the critical concentration for BtubA/B assembly (Fig. 3B and Table 1). To deconvolve the effects of KCl and polymer concentration on bundling, we compared BtubA/B assembly kinetics in conditions that produce a similar mass of bMTs (21) (Fig. 3C). Polymerization curves shows that at a similar bMT concentration the increment of KCl increases the light scattering intensity, indicating that filament bundling is mainly influenced by KCl over bMT concentration.

**Fig. 3.**
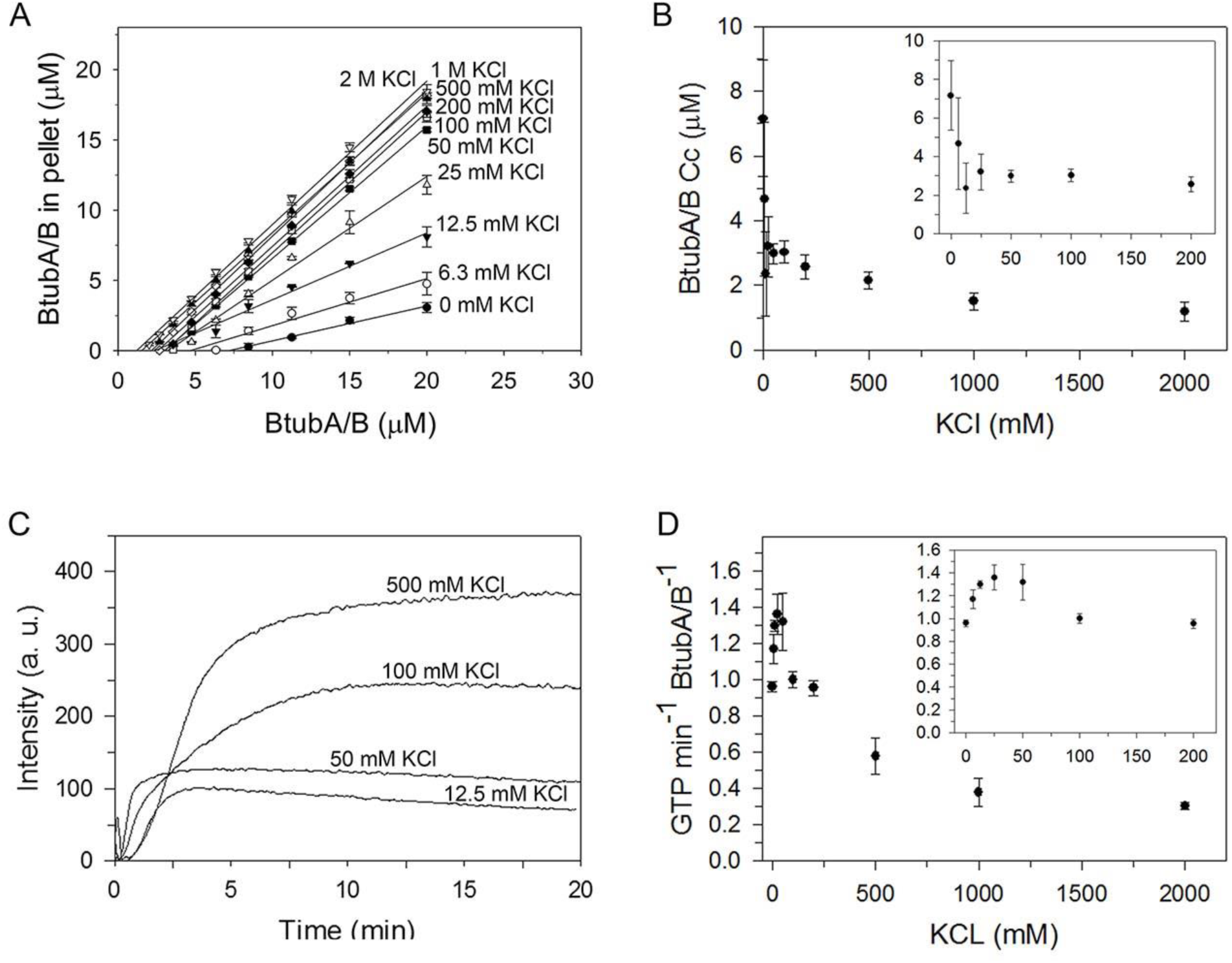
The increase of KCl increases the mass of assembled bMTs and reduces GTP hydrolysis. (A) Different BtubA/B concentrations were polymerized at different KCl concentrations by addition of 1 mM GTP, and centrifuged at 25°C at 100,000 x g. The amount of BtubA/B pelleted was plotted against total BtubA/B concentration. The x-intercept of the linear fit corresponds to the BtubA/B critical concentration. (B) Critical concentration of BtubA/B polymerization versus KCl concentration. Inset correspond to an expanded view at lower KCL concentrations. (C) Comparison of four BtubA/B light scattering polymerization curves (data from Fig. 2 and Fig. S2 in the supplemental material) based on the bMT concentration determined by High-speed pelleting (Fig. 3A). Polymerization conditions were: 8.4 μm BtubA/B in 12.5 mM KCl (3.2 μm bMTs), 6.3 μm BtubA/B in 50 mM KCl (3.2 μm bMTs), 4.7 μm BtubA/B in 100 mM KCl (1.6 μm bMTs),and 3.6 μM BtubA/B in 500 mM KCl (1.3 μM bMTs). Polymerization kinetics in 12.5 mM KCl exhibited sigmoidal behavior and the lowest maximum intensity between the conditions. However, increasing KCl to 50 mM KCl increased the light scattering intensity. The increase in signal intensity was greater in the transition to 100 mM KCl, where the trace exhibited an elongation phase composed by two steps as previously described (Fig. 2C). At 500 mM KCl, two steps cannot be resolved in the elongation phase, and depolymerization is slower. This indicates that bMT stabilization by bundling is principally influenced by KCl, rather than protein concentration. (D) GTPase activity of BtubA/B *versus*KCl concentration. Inset: expanded view of GTPase activity at low salt.

### Effect of KCl on GTPase activity of BtubA/B

Because the dynamic assembly and disassembly of eukaryotic microtubules and bacterial FtsZ filaments are regulated by GTP binding, hydrolysis, and exchange, we analyzed BtubA/B GTPase activity at different KCl concentrations (Fig. 3D and Table 1). In the absence of KCl, the GTPase activity was 0.96 ± 0.03 mol GTP per minute per mole of BtubA/B, which increased slightly to 1.36 ± 0.11 mol GTP per minute per mole of BtubA/B at 25 mM KCl. However, at KCl concentrations above 50 mM, the GTPase activity decreased continuously to 0.3 ± 0.02 mol GTP per minute per mole of BtubA/B in 2 M KCl.

### BtubA/B polymerization is kinetically polarized

Like eukaryotic microtubules, the BtubA/B polymer is thought to be structurally polar (3), but this does not necessarily imply that the kinetics of assembly are also polar. For example, the structurally polar actin homolog ParM (22) elongates symmetrically (23, 24). To test the hypothesis that bMTs exhibit kinetic polarity, we studied bMT assembly by fluorescent filament imaging (23) using BtubA/B seeds labeled with TAMRA (red) and soluble BtubA/B labeled with Alexa Fluor 488 (green) at low salt concentration to avoid the confounding effects of bundling. Most of the observed filaments (90%) had one red end and one green end (Fig. 4), indicating that, as for eukaryotic microtubules, the growth of bMTs is kinetically polar. Polar bMTs were clearly observed from 12.5 mM KCl up to 100 mM KCl (see Fig. S3 in the supplemental material). At higher KCl concentrations, we observed thicker yellow polymers, which we interpret as mixed-polarity bundles of bMTs. Following eukaryotic microtubule nomenclature, we refer to the fast growing end as the bMT plus end (+) and to the slow growing end as the bMT minus end (−).

**Fig. 4.**
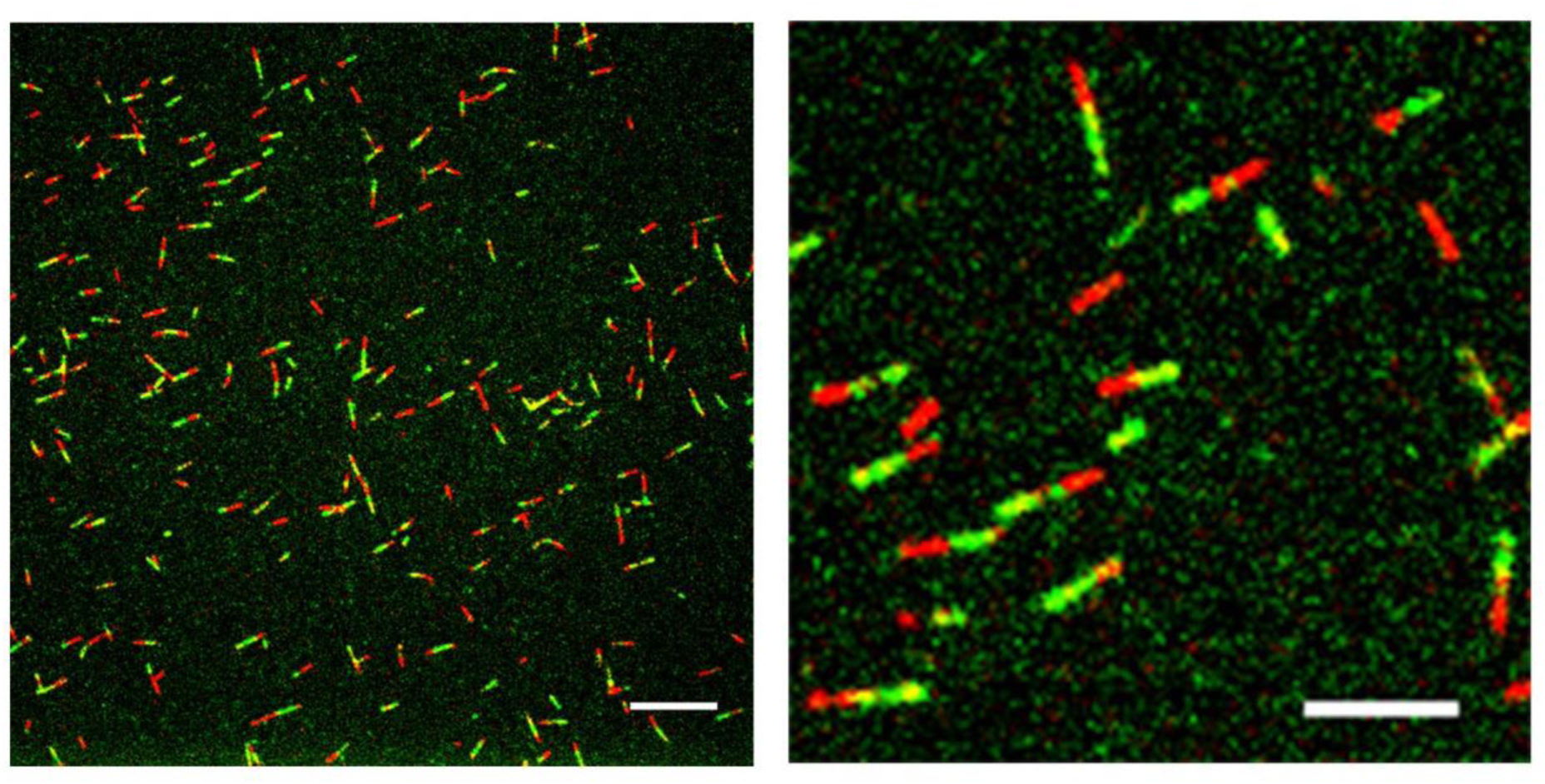
bMT assembly is kinetically polar. Kinetic polarity assay of bMTs (see Materials and Methods). TAMRA-labeled BtubA/B (red) was polymerized with 1 mMΜMPCPP in 25 mM KCl. After 2 min, Alexa Fluor 488-labeled BtubA/B (green) was added. Right image correspond a magnification of the left image. (Scale bar: 10 μm in left image; 5 μm in panel in right image.)

### bMT bundle and filament assembly in high salt

To gain insights into bMTs assembly dynamics, we monitored the assembly dynamics of fluorescently labeled bMTs growing from seeds stabilized with ΜMPCPP (a slowly hydrolyzable GTP analogue) at 100 mM KCl using confocal fluorescence microscopy. Consistent with our bulk polymerization results, in 100 mM KCl and GTP, bMTs growing at 3.7 ± 0.2 μM min^-1^ (n= 8), formed long and interconnected bMT bundles (Fig. 5A and Movie S1 in the supplemental material). bMT bundling occurs through the interaction of parallel or antiparallel individual growing bMTs, forming thin bundles, which in turn interact together (zippering) to form thicker and longer bundles (as observed by an increase in fluorescence intensity) (Fig. 5B and Movie S2). Some ΜMPCPP-stabilized bMT seeds also elongated as a bipolar bundle (Fig. S4), confirming that bundles are composed of antiparallel bMTs. Furthermore, non-uniform fluorescence intensity along the bMT bundles, especially the movement of bright spots along bundles, suggests that bMTs may still elongate and exhibit dynamics while associated with a bundle (Fig. 5C and Movie S3)

**Fig. 5.**
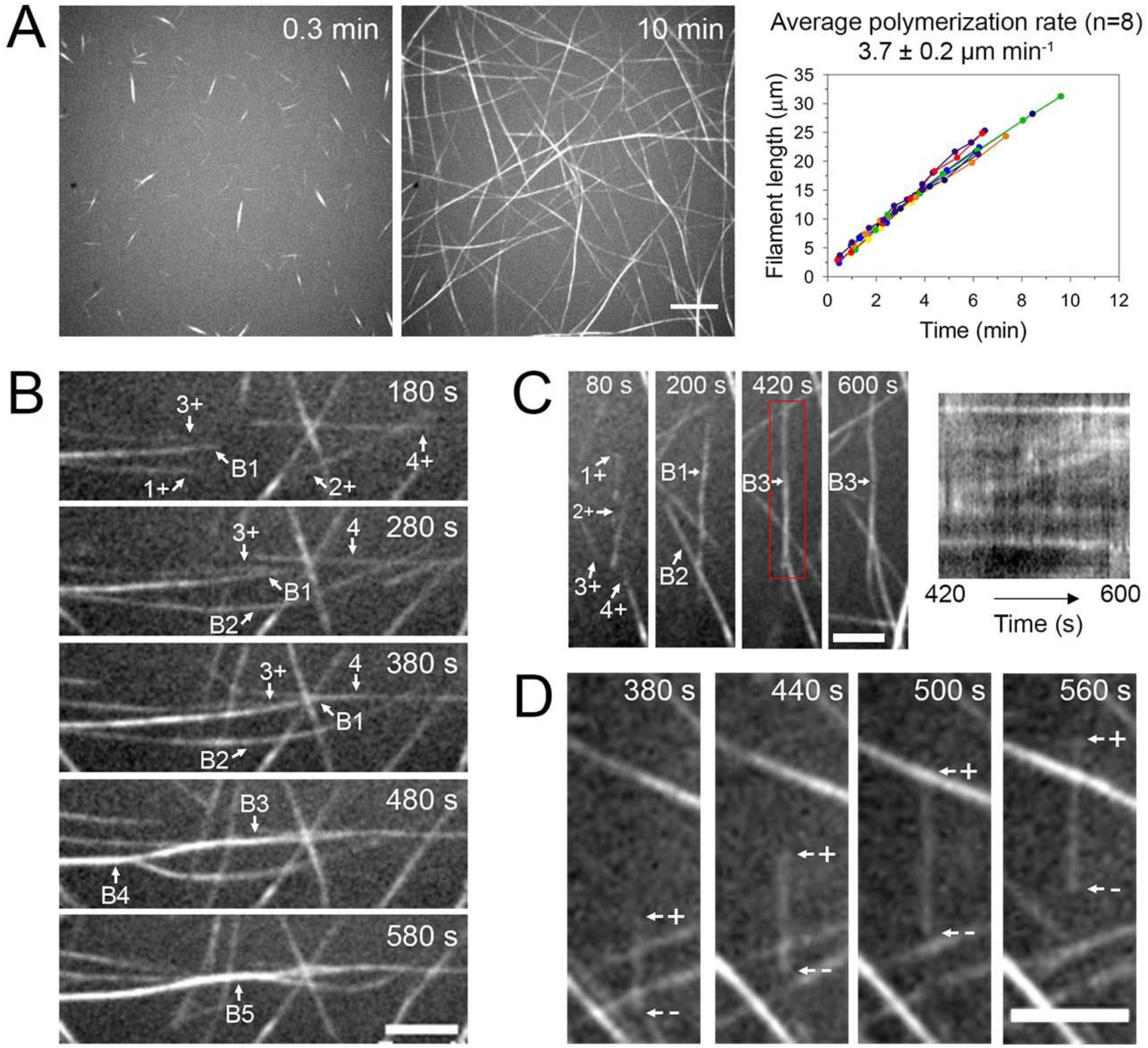
Visualization of BtubA/B polymerization and bMT bundling at high KClconcentration. (A) Time course of bMT polymerization (3 μm BtubA/B, 30% TAMRA-labeled) from ΜMPCPP-stabilized seeds (brighter labeling) in 50 mM HEPES (pH 7), 5 mM MgCl_2_, 100 mM KCl, and 1 mM GTP. Right plot: filament length *versus* time for eight individual bMTs. (B) Two bMTs growing in opposite directions (“1” and “2”, where “+” denotes the growing end of the bMT) interact end-to-end, forming a bundle (B2). Two single bMTs (“3+” and “4+”) and a bundle (B1) interact longitudinally to form a thicker bundle (B3). At 480 s and 580 s bundles B2 and B3 zipper together (B4 and B5). (C) Dynamics of bMT bundles. Left: a bundle (B1) formed by the interaction of two parallel growing bMTs (“1+” and “2+”) interacts with a bundle (B2) formed by two antiparallel growing bMTs (“3+” and “4+”). Right: kymograph between 420-600 s of the bMT bundle dynamics (the region analyzed in the kymograph is enclosed by the red rectangle). Non-uniform fluorescence along the bundle suggests that bMTs polymerize in the bundle. (D) bMTtreadmilling. The rate of polymerization of the bMT growing end (“+”) is similar to the depolymerization rate at the other end (“-“). (Scale bar: 10 μm in panel (A); 5μm in panel (B), (C) and (D).)

Interestingly, we observed bMT treadmilling, consisting simultaneous growth at the plus end and subunit loss at the minus end (Fig. 5D and Movie S4; Fig. S5 and Movie S5), in fragments whose minus ends were not stabilized by a seed. These bMT fragments indicate that breakage events do occur in these conditions, and that the critical concentration must be different between their plus and minus ends.

### bMTs in low salt exhibit steady growth punctuated by stochastic fragmentation

In order to explore the dynamics of single filaments in more detail without the confounding effects of bundling, we visualized the assembly of TAMRA-labeled BtubA/B in 12.5 mM KCl and GTP (Fig. 6A and Movie S6 in the supplemental material). We observed single bMTs growing from seeds in one direction, and two or more filaments growing from seeds in opposite directions, with an average rate of 2.4 ± 0.3 μm min^−1^ (n= 7).

**Fig. 6.**
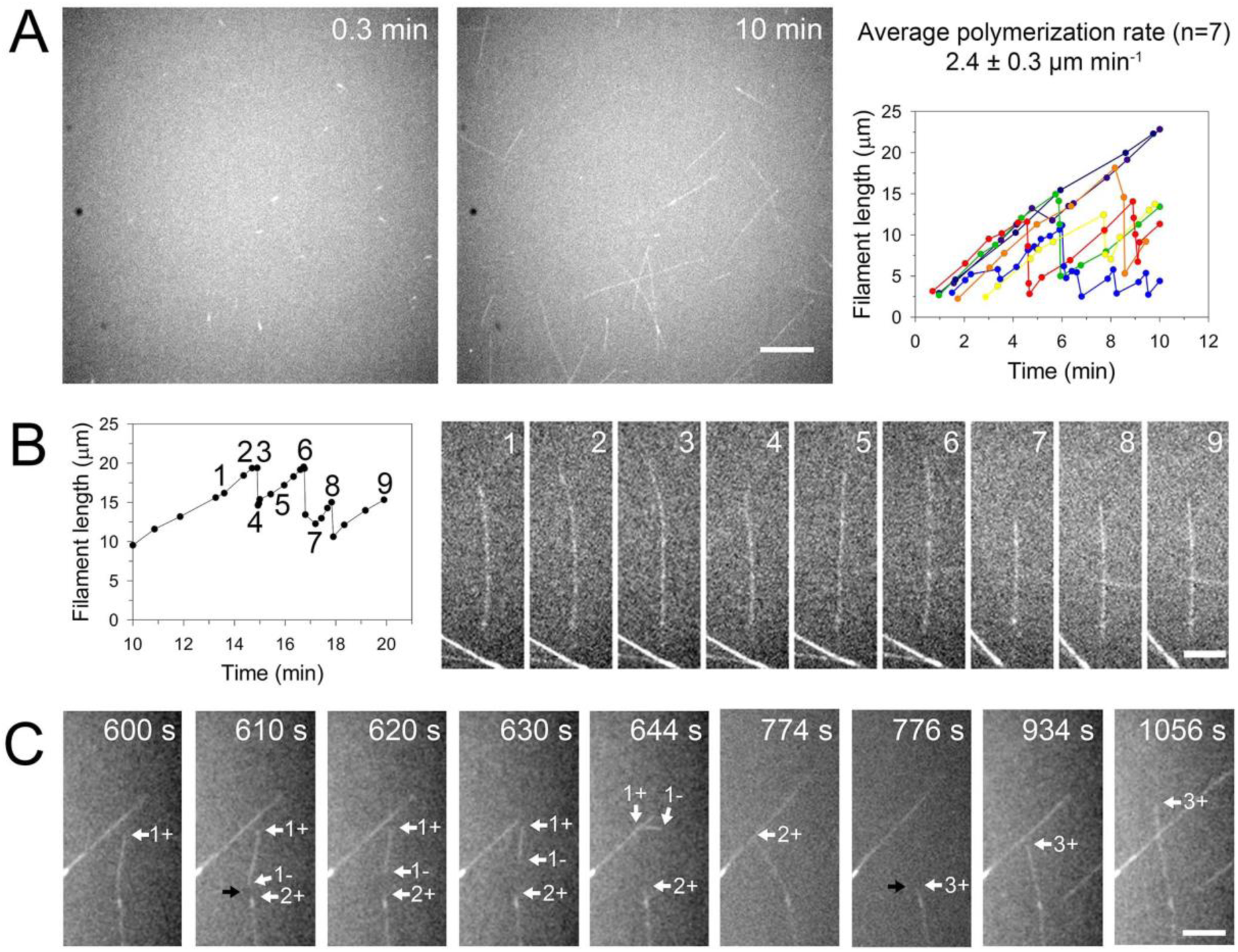
Visualization of BtubA/B polymerization at low KCl concentration. Bacterial microtubules alternate between steady growth and fragmentation. (A) Time-lapse images of 3 μm BtubA/B (30% TAMRA-labeled) assembly from seeds in 12.5 mM KCl and 1 mM GTP. Right: filament length versus time for seven single bMTs growing from ΜMPCPP-stabilized seeds. (B) Left: change in length as a function of time of the single bacterial microtubule. Right: Dynamically unstable bMT elongating from a seed at 2.2 ± 0.2 μm min^-1^. (C) Spontaneous breakage (black arrow) of a growing bMT (“1+” where “+” denotes the growing end of the bMT) generates a fragment that remains bound to the surface. The new plus end (“2+”) resumes growth and the new minus end of the bMTfragment (“1-“ where“-“ denotes the shrinking end) starts depolymerizing faster than the plus end of the bMTfragment (“1+”). 776 s: spontaneous breakage (black arrow) generates a new growing plus end (“3+”). (Scale bar: 10 μm in panel (A) and (C); 5 μm in panel (B).)

We observed that bMTs suddenly switch from steady growth to a shortening phase (Fig. 4A and B, and Movie S6 and S7). Some bMTs fragmented completely (exposing the seed end), while other bMTs reduced in length. In both cases, the elongation phase from the plus end resumed at a similar rate as before. The elongation time until the first breakage event varied among the filaments, indicating stochastic switching between elongation and shortening (23). In most cases, bMT fragments resulting from breakage were not observable, indicating that fragments quickly detached from the surface or that the depolymerization from their new minus end, no longer stabilized by GMPCPP, was so fast that we were unable to visualize it. However, we also occasionally observed bMT fragments bound to the surface after bMT breakage, which depolymerized from their new minus ends at a faster rate than the growing plus ends, causing the disassembly of the bMT fragment (Fig. 6C). As expected, the increase of BtubA/B concentration from 3 μm to 5 μm produced longer and faster-growing filaments (Fig. S6A and Movie S8 in the supplemental material), and also increased the lifetime of bMT fragments without stabilized minus ends (Fig. S6B in the supplemental material). We also observed bMT breakage caused by a growing bMT end running into a perpendicularly aligned bMT (Fig.S6C in the supplemental material).

The dynamic instability model of eukaryotic microtubules predicts that the stochastic shrinkage is dependent on GTP hydrolysis in the MT wall (10–12, 15). We determined the effect of GTP hydrolysis inhibition on bMT dynamics by visualizing BtubA/B polymerization in 12.5 mM KCl and in the presence of the non-hydrolyzable GTP analog MPCPP. We observed a large number of short bMT bundles about 10 μm in length, and between 5 and 10 minutes time points, bMT bundles were not dynamic (Fig. 7A). The bidirectional polymerization of bMTs from seeds at low KCl concentration (Fig. 6A and Fig. S6A in the supplemental material) indicates that seeds can form bundles composed of antiparallel bMTs even in low salt. In most experiments, the ΜMPCPP-stabilized seed mixture was centrifuged to remove excess GMPCPP. The high concentration in the seed pellet may induce bundling. To reduce the formation of bundled seeds, we synthetized the seeds without centrifugation, added the seed solution directly to the reaction chamber, and washed out the GMPCPP while the seeds were attached to the surface. We then flowed in 5 μm BtubA/B in 12.5 mM KCl and 1 mM GTP. The growing bMTs formed long and stable bundles without dynamic rearrangement over time (Fig. 7B and Movie S9 in the supplemental material). We interpret this data as evidence that even trace amounts of GMPCPP remaining in the reaction chamber can stabilize the bMTs against breakage.

**Fig.7.**
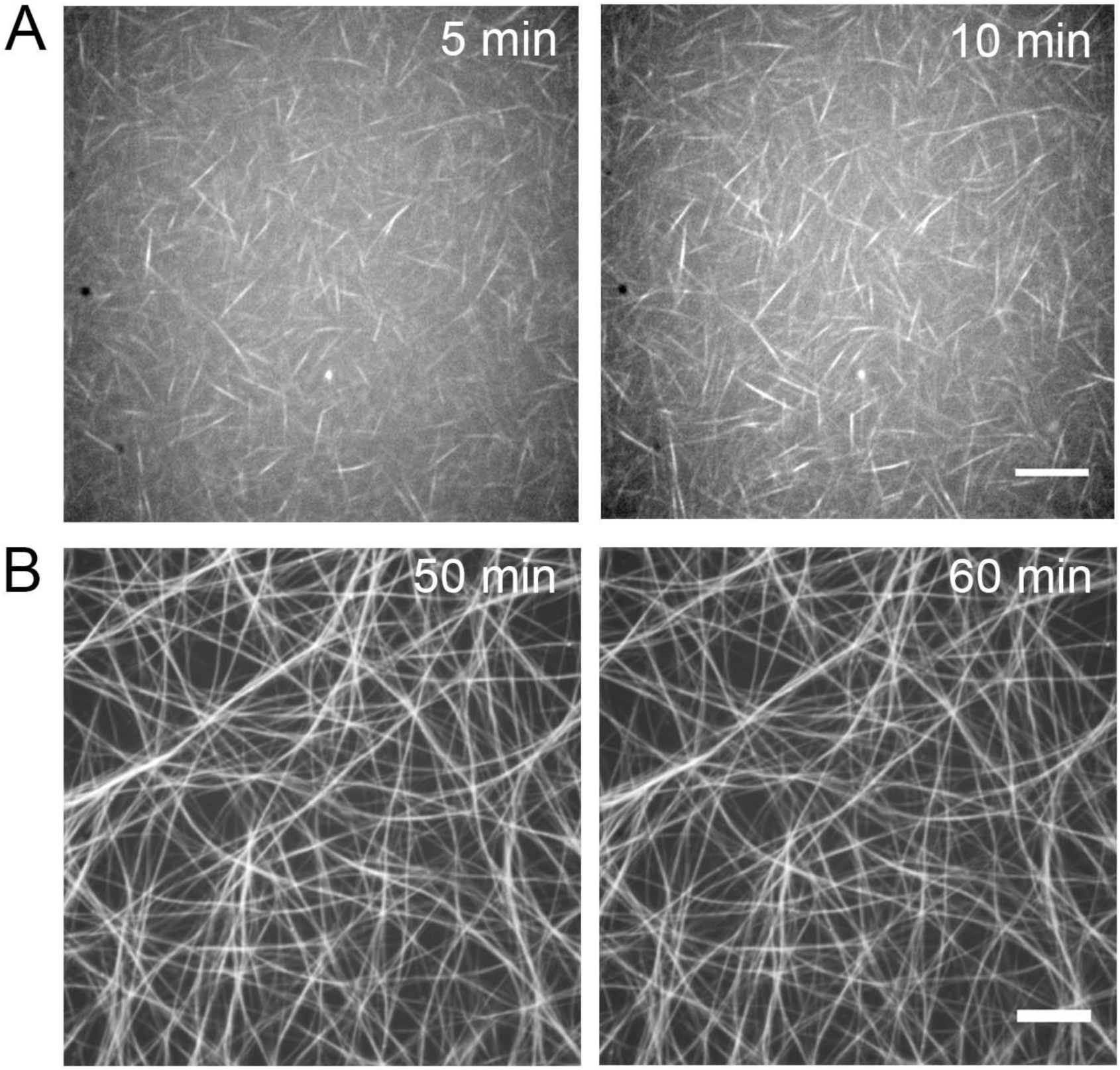
BtubA/B polymerization with the slowly hydrolyzable analog GMPCPP. (A) 5 μM mBtubA/B (30% TAMRA-labeled) polymerized in the presence of 1 mMΜMPCPP and 12.5 mM KCl. Seeds are brighter and denser than in polymerization with GTP (Fig. S6A), indicating that they contain several bundled filaments. (B) 5 μmBtubA/B (30% TAMRA-labeled BtubA/B) polymerized in the presence of 12.5 mM KCl by adding 1 mM GTP, and trace GMPCPP. bMTs form long and interconnected bundles that remain stable even at 60 min after GTP addition. Transitions between bMT growth and fragmentation were not observed. (Scale bar: 10 μm in panel (A) and (B).)

## Discussion

In this study, we have characterized the biochemical properties, assembly polarity, and dynamics of bacterial microtubules. We determined that BtubA/B assembly is kinetically polar and its assembly dynamics are affected by KCl concentration. At low KCl concentration, single bMTs growth and shrink via stochastic filament fragmentation. The increase of KCl concentration reduce the critical concentration of BtubA/B assembly and induce bMT bundling.

We imaged the products of BtubA/B polymerization in a range of salt conditions using both fluorescence and electron microscopy (Fig. 1). At low salt, we observed single bMTs, which, under negative stain, appear as two light filaments flanking a dark center, which are consistent with either paired protofilaments or the light tubule walls flanking a dark tubule lumen. This appearance of the tubules is similar to previous observations of BtubA/B polymerized *in vitro* and imaged with negative stain electron microscopy (2, 4, 7). These results were initially interpreted as evidence that BtubA/B filaments are composed of protofilament doublets, but subsequent electron cryotomography of BtubAB assembled in bacteria showed that bMTs are hollow tubules composed of 5 protofilaments (1). Such a 5-protofilament tubule structure has a similar overall diameter and appearance when viewed in projection, as in negative stain electron microscopy. There may therefore be no major structural difference between bMTs assembled *in vitro* and bMTs in bacteria. Using light scattering (Fig. 2 and Fig. S2 in the supplemental material), we determined that below 25 mM KCl, bMTs follow a nucleation-elongation mechanism of assembly. At 50 mM or higher KCl concentrations, bMTs self-associate into bundles, altering the shape of the kinetic trace, and reducing bMT depolymerization rate compared to low KCl concentration, indicating that bundling stabilizes the filaments against depolymerization. The comparison between polymerization traces with similar polymer amount but different KCl concentrations indicates that bMT stabilization by bundling is principally influenced by KCl, rather than protein concentration (Fig. 3C).

The reduction of the critical concentration of BtubA/B assembly increases the mass of assembled bMTs (Fig. 3A). KCl may decrease the critical concentration by inducing bundling, which stabilizes heterodimer interactions in filaments. However, the reduction of critical concentration at low salt where there is no bundling (Fig. 2E and Fig. 3B) suggests that KCl has also a stabilizing effect on the BtubA/B interaction along a single bMT. This KCl effect could be favoring both intra-and inter-heterodimer interaction as suggested by the increase of the size of the BtubA/B polymerization nucleus with increasing KCl concentration (Fig. 2F). The increase of the nucleus size from 0 (n=~0.5), 3.1 (n=~1), and 6.3 mM KCl (n=~1.4) indicates that KCl favors heterodimer formation or intra-heterodimer interaction, a step that is essential for BtubA/B polymerization (REF). At 12.5 mM and 25 mM KCl, nucleus is composed by two BtubA/B heterodimers suggesting that KCl also increases the inter-heterodimer interaction. In a complementary approach, we measured effective total GTP hydrolysis rate to increase at low salt and to decrease with salt-induced bundling from 50 mM KCl (Fig. 3D and Table 1). The increase of GTPase activity from 0 to 25 mM KCl could be explained by an increase of BtubA/B competent to polymerize (and to hydrolyze GTP) due to the reduction of the critical concentration. On the other hand, the reduction of GTPase activity from 50 mM KCl correlates with the onset of bMT bundling (Fig. 1B and Fig. S2D in the supplemental material) and it could be explained by a reduction in free (unpolymerized) BtubA/B heterodimer due to tubule stabilization by bundling. bMT bundling therefore appears to be an important modulator of bMT dynamics, which may have implications for their function in either *Prosthecobacter* or in the ancient eukaryotic ancestor in which the ancestral microtubule existed.

We visualized the dynamics of fluorescently labeled bMTs using confocal fluorescence microscopy and determined that, like eukaryotic MTs, bMTs are kinetically polar polymers (Fig. 4). In the presence of GTP, bMTs exhibit complex dynamics including continuous treadmilling (Fig. 5D) and stochastic switching between growth and fragmentation (Fig. 6B). Treadmilling has also been observed in the plasmid-encoded bacterial tubulin TubZ (25), FtsZ filaments (26) as well as in eukaryotic MTs. bMT treadmilling indicates that GDP/GTP exchange can only occur in the free heterodimers, because it occurs when monomers at the plus end are GTP-bound and monomers at the minus end are GDP-bound. Such a hydrolysis state gradient is inconsistent with exchange in the bMT wall. The stochastic switching between growth and disassembly is similar to the dynamic instability seen in eukaryotic MTs as well as in filaments formed by the distantly related bacteriophage phage tubulin PhuZ (27) and the actin-related protein ParM (19, 23). These dynamic behaviors are suppressed by the slowly hydrolysable GTP analogue GMPCPP (Fig. 7), indicating that GTP hydrolysis is necessary for bMT destabilization and disassembly, as for eukaryotic MTs.

In eukaryotic MTs, loss of the stabilizing cap of GTP-bound monomers at the plus end leads to filament depolymerization and the gain of a GTP cap rescues polymerization (10–12). With the imaging modality and fluorescent labeling used in this study, it was not possible to observe the fast continuous depolymerization at the new plus end created by a break, which would unequivocally confirm dynamic instability. However, bMT fragmentation is consistent with a GTP hydrolysis-induced loss of filament stability and we have observed rescue of polymerization after fragmentation (Fig. 6B). The lower number of protofilaments in bMTs may render them more prone than eukaryotic MTs to fragmentation. Fast disassembly may then occur as a combination of fragmentation and fast depolymerization at the new plus end. We conclude that at low KCl concentration, single bMTs are mechanically unstable filaments with transitions between steady growth and disassembly, consisting of bMT fragmentation, processes that are both modulated by GTP hydrolysis state.

Our experiments in a range of buffer conditions show that bMTs differ from eukaryotic MTs in their strong tendency to assemble into irregular and apolar bundles in high salt concentration. Bundling does not require any other proteins. Pre-existing single bMTs can zipper together into bundles and keep growing (Fig. 5C) and on rare occasions treadmill while associated with a bundle (Fig. S5>). bMTs stabilized by GMPCPP were observed to bundle even in low salt (12.5 mM KCl, Fig. 7), possibly due to the sum of many weak interactions along their lengths. An attractive but non-stereospecific interaction, such as hydrophobic interaction or van der Walls forces between bMTs may be responsible for the bundling, and for short filaments, salt is needed to screen the charge on bMTs. Crowding agents also enhance BtubA/B polymerization and induce the bundling of bMTs (7). We did not observe breakage of bundled bMTs, suggesting that bundling mechanically stabilizes the bMT.

The function of bMTs *in vivo* is still unknown. In *Prosthecobacter,* single bMTs of 0.2−1.2 μM in length, or bundles composed by 2, 3, or 4 bMTs, are located in or near the base of the cellular extension called prostheca(1). bMTs or their bundles may support the elongated shape of the prostheca (4, 7), but so far, evidence suggests that bMTs are not determinants of the *Prosthecobacter* cell morphology (28). Based on our *in vitro* observations, we propose that *in vivo*, bMT bundles are composed of antiparallel filaments and bMT bundling should be favored as the predominant supramolecular arrangement of bMTs, (assuming that *Prosthecobacther* cytoplasm is similar to *E. coli* cytoplasm (0.1 −1.0 M K^+^(29), 1−5 mM Mg^+2^(30), pH≈ 7.7 (31)). Due to the lack of genetics tools in *Prosthecobacter,* the controlled expression and visualization of fluorescently tagged BtubA/B in *E. coli* would be useful steps to determine the dynamics and function of bMTs in a cell environment. Ultimately, bMT function in *Prosthecobacter* will also need to be understood in the context of bMT-binding proteins that potentially regulate their dynamics *in vivo,* such as the protein product of the *bklc* gene (6).

In addition to their unknown function in *Prosthecobacter*, bMTs are of ultimate interest as an evolutionary link between eukaryotic microtubules and the ancient microtubule from which the BtubA/B genes are derived. BtubA/B has more sequence and structure in common with eukaryotic tubulins than any other bacterial tubulin, and the presence of both *btubA* and *btubB* genes suggests that the horizontal transfer event that is thought to have occurred not long after the initial duplication event that created the ancestors of the A/B and α/β homolog pairs (9). The more evolutionarily distant phage tubulin PhuZ has been demonstrated to exhibit dynamic instability (27) and to segregate DNA (32). We cannot exclude the possibility that bMT dynamics evolved after the horizontal gene transfer to *Prosthecobacter,* and any similarities between the dynamics of bMTs, phage tubulin filaments and eukaryotic MTs are the result of convergent evolution. In either situation, however, BtubA/B provides a simple model of bacterially expressed tubulins that could be used to study the structural determinants of microtubule dynamics.

Further work on the structural changes in the BtubA/B dimer in response to GTP hydrolysis and polymerization, as well as characterization of the *bklc* gene product will illuminate the extent to which BtubA/B and the ancient tubulin ancestor could have functioned in DNA segregation or other cellular functions.

## Materials and Methods

### BtubA/B purification

A vector containing the *Prosthecobacter dejongeii btubA* and *btubB* genes under control of a T7 promoter (kindly provided by D. Schlieper and J. Löwe) was transformed into *E. coli* BL21(DE3) cells. A single *E. coli*BL21(DE3) colony was grown at 37°C to an optical density of 0.6 at 600 nm, and BtubA/B overexpression was induced by adding 0.3 mM IPTG to the cell culture. After 4 hours, the cell culture was centrifuged at 5,000 x g for 35 min and the cell pellet was suspended with ice-cold TEN buffer (50 mM Tris-HCl (pH 8), 1 mM EDTA, 100 mM NaCl). This procedure was repeated twice to wash away LB medium, and then was repeated 2 times using 50 mM HEPES-KOH (pH 7) buffer. The cell pellet was frozen at −80°C for later use. The pellet was thawed on ice and suspended in 4 volumes of cold 50 mM HEPES-KOH (pH 7) and 1 mMΜMSF. The suspension mixture was sonicated on ice seven times with twenty second bursts at 7.0 W with a Misonix 3000 sonicator. The lysate was centrifuged at 100,000 x g for 1 h at 4°C, the supernatant was recovered and the pellet was discarded. The supernatant was warmed to 25 °C, its volume was measured and 5 mM MgCl_2_ (2 M stock solution) and 0.5 M KCl (solid, to minimize the increase in volume) were added. The solution was mixed gently and it was incubated at 25 °C for 10 min. The first BtubA/B polymerization/depolymerization cycle was started by adding 2 mM GTP. The solution was gently mixed and centrifuged immediately at 100,000 x g for 30 min at 25 °C. The supernatant was discarded and the pellet was solubilized by pipetting with 3-4 volumes of ice-cold 50 mM HEPES-KOH (pH 7) buffer, then was incubated in ice for 30 min (low temperature reduces BtubA/B polymerization, as with eukaryotic MTs), with occasional pipetting, and was ultracentrifuged at 100,000 x g for 30 min at 4°C. The pellet was discarded. This completed one polymerization/depolymerization cycle. The supernatant was warmed again at 25 °C and a total of three cycles of polymerization/depolymerization were completed using the same conditions. The last solubilized pellet was dialyzed twice against 1000 volumes of 50 mM HEPES-KOH (pH 7) buffer, then the protein solution was ultracentrifuged at 100,000 x g for 15 min at 4°C to remove aggregates, and the protein in the supernatant was stored in aliquots at −80 °C.

### Quantification of BtubA/B concentration

BtubA/B batch stock concentration was determined by the BCA assay and using BtubA/B of known concentration as a standard. BtubA/B standard concentration was quantified by measuring its absorbance at 280 nm after nucleotide exchange: an aliquot of BtubA/B was incubated with 2 mM GTP and 5 mM MgCl_2_ for 1 h in ice to exchange any GDP bound to BtubA/B. The free nucleotide was removed using a Bio-Gel P-6 polyacrylamide gel chromatography column (Bio-Rad), previously equilibrated with polymerization buffer (50 mM HEPES-KOH (pH 7), 5 mM MgCl_2_). The eluent was ultracentrifuged at 100,000 x g for 15 min at 4°C to remove aggregates and BtubA/B polymers. A 1:100 dilution of the supernatant (BtubA/B bound to GTP) in polymerization buffer was performed to measure its absorbance at 280 nm using an extinction coefficient of 103,754.2 M^-1^ cm^-1^ (BtubA, 47,900 M^-1^ cm^-1^;BtubB, 39,880 M^-1^ cm^-1^; 2 GTP (7987.1 M^-1^ cm^-1^ each). For the light scattering, high-speed pelleting and GTP hydrolysis assays, 20 μmBtubA/B in different KCl polymerization buffers was prepared from 400 μmBtubA/B stock, resulting in a KCl and MgCl_2_ concentration decrease of 5% from the nominal value. The 20 μmBtubA/B solution was serially diluted in 0.75-fold increments (20, 15, 11.3, 8.4, 6.3, 4.8, 3.6, 2.7, 2 and 1.5 μmBtubA/B) using the same KCl polymerization buffer.

### Fluorescent Labeling of BtubA/B

200 μmBtubA/B in 50 mM HEPES (pH 7) was fluorescently labeled by adding a 10-fold molar excess of TAMRA (5-(and-6)- carboxytetramethylrhodamine, succinimidyl ester, mixed isomers, Molecular Probes) or Alexa Fluor 488 (Molecular Probes), for 1 h on ice with occasional stirring. Labeled BtubA/B was separated from the free dye by Bio-Gel P-6 polyacrylamide gel chromatography columns (Bio-Rad). Labeled BtubA/B was warmed to room temperature and 200 mM KCl and 5 mM MgCl_2_ were added to it. Labeled and functional BtubA/B was selected by the addition of 1 mM GTP and the solution was immediately centrifuged at 100,000 x g for 15 min at 25°C. The pellet was suspended in 50 mM HEPES-KOH (pH 7) and centrifuged at 100,000 x g for 15 min at 4°C. The supernatant was dialyzed against 50 mM HEPES-KOH (pH 7) for 4 h and then was stored in aliquots at -80°C. Final labeling stoichiometry was 1-2 dyes per BtubA/B heterodimer.

### Electron microscopy

5 μm BtubA/B in polymerization buffer (50 mM HEPES-KOH (pH 7), 5 mM MgCl_2_) with different KCl concentrations was polymerized by adding 1 mM GTP at 25°C. After 5 min, 4 μL of the reaction mixture was transferred to glow-discharged 200- mesh carbon-Formvar-coated cooper grids at 25°C. The grids were washed with three drops of the same KCl polymerization buffer and were negatively stained with 0.75% uranyl formate. The sample images were obtained at 25°C with a Tecnai T12 microscope using an acceleration voltage of 120 kV and a magnification of 52,000X. Images were recorded with a Gatan 4k × 4k charge-coupled device camera.

### Light scattering

BtubA/B polymerization kinetics was measured by 90° light scattering on a Perkin-Elmer spectrofluorimeter LS-50B, using an excitation and emission wavelength of 350 nm, 5 nm excitation and emission bandwidths, and a transmission filter of 4%. Each BtubA/B solution was loaded in a quartz cuvette with a 1 cm path length and the light scattering of the solution was measured for 5 min at 25°C to establish the baseline. Polymerization was initiated by adding 1 mM GTP to the cuvette, which was gently homogenized by inversion (3 times). The light scattering intensity was measured starting 15 s after the addition of nucleotide. The steady state BtubA/B concentration was determined from the difference between the maximum intensity value and the average intensity of the 5 min baseline of each kinetic trace. This net intensity was plotted as a function of the initial BtubA/B concentration. The x-axis intercept of the linear fit to the plot corresponds to the critical concentration of unpolymerized BtubA/B.

### High-speed pelleting

For the pelleting assays, reactions started by adding 1 mM GTP to the BtubA/B solutions and were immediately centrifuged at 100,000 x g in a TLA 100.4 rotor for 15 min at 25°C. Precipitates were suspended in the starting volume of the reaction without KCl in the polymerization buffer, and BtubA/B was quantified by the BCA assay, using the serial dilution of BtubA/B as a calibration curve. The critical concentration for BtubA/B assembly was determined by plotting BtubA/B in the pellet as a function of the initial BtubA/B concentration. The x-axis intercept of the linear fit to the plot corresponds to the steady state concentration of unpolymerized BtubA/B.

### GTP hydrolysis assay

GTPase activity (phosphate release (Pi)) was measured by the malachite green colorimetric assay (33). The reaction was started by adding 1 mM GTP to the BtubA/B solutions. At 0 and 5 minutes, 50 μL aliquots were added to tubes containing 350 μL nanopure water and 400 μL perchloric acid, mixed and placed in ice to stop GTP hydrolysis. After 30 min, 200 μL of the dye solution was added to 800 μL sample, the sample was incubated for 10 min at 25°C and absorbance was measured at 630 nm. The standard curve was constructed with monobasic potassium phosphate dissolved in nanopure water, and had a linear range between 0.5 and 10nmol of inorganic phosphate. The rate of GTP hydrolysis (Pi [μm] min^-1^) was calculated from the difference between the measurement at 5 minutes and time 0. We plotted the rates of GTP hydrolysis as a function of BtubA/B concentration. The slope of the line fit corresponds to BtubA/B GTPase activity.

### Kinetic polarity assay of bMT assembly

20 μL of 5 μm BtubA/B (30% TAMRA-labeled) was assembled by the addition of 0.5 mMΜMPCPP (50 mM HEPES-KOH (pH 7), 5 mM MgCl_2_, 0-500 mM KCl). After 2 min, 20 μL of 5 μm BtubA/B labeled with Alexa Fluor 488 (30% Alexa Fluor 488-labeled) was added to the reaction mixture. After 2 min, 1 μL of the reaction was diluted in 200 μL of polymerization buffer and 4 μL of the dilution was added to a poly-L-lysine-coated coverslip and visualized by confocal microscopy.

### Confocal Microscopy

For confocal microscopy visualization of in bulk BtubA/B assembly, 5 μm BtubA/B (15% TAMRA-labeled) in polymerization buffer (50 mM HEPES-KOH (pH 7), 5 mM MgCl_2_) with different KCl concentrations (50, 100, and 1,000 mM KCl) was polymerized by adding 1 mM GTP. After 5 min, the solutions were transferred to acid-washed and poly-L-lysine coated cover slips before visualization. Individual bMTs growing from seeds were visualized by confocal microscopy in flow chambers made of acid-washed cover slips. Cover slips were kept overnight in 1 M HCl at 60°C, then sonicated for 30 min in MilliQ (Millipore) distilled deionized water (twice) and in a ramp of 50%, 75%, and 95% ethanol washes, then stored in 95% ethanol. Cover slips were rinsed thoroughly with MilliQ water and dried with a nitrogen stream before use. The assembly assay was adapted from the method described by Brouhard et al. (34) with minor modifications. Flow chambers were incubated with anti-rhodamine antibody (Invitrogen) diluted 1:100 in polymerization buffer at room temperature for 15 min. The flow chamber was washed with polymerization buffer followed by two washes of blocking solution (1% pluronic acid 127 (Sigma) in polymerization buffer) for 10 min. BtubA/B seeds diluted 1:100 in blocking buffer were added to the flow chambers and were incubated for 5 min at room temperature. The flow channels were washed with blocking buffer and the BtubA/B assembly reaction was added to the flow channels. The BtubA/B assembly reaction was supplemented with glucose oxidase oxygen scavenger system consisting of 250 μg/mL glucose oxidase (Sigma-Aldrich), 4.5 mg/mL glucose, and 30 μg/mL catalase (Roche). The BtubA/B seeds were synthetized by polymerizing 5 μm BtubA/B (40% TAMRA-labeled) in polymerization buffer with 12.5 or 100 mM KCl by adding 0.5 mMΜMPCPP for 5 min at room temperature. The solution was centrifuged at 100,000 x g for 15 min, the supernatant was discarded and the pellet was suspended in polymerization buffer, followed by a second centrifugation and suspension in polymerization buffer.

Samples were imaged with a Carl Zeiss (Oberkochen, Germany) Axio Observer Z1 inverted microscope with a Zeiss 63x, 1.4 NA Plan Apochromat oil immersion objective, a Yokogawa (Sugar Land, TX) CSU 10 confocal scanner unit, and a Photometrics (Tucson, AZ) Cascade II 512 CCD camera. For confocal illumination, Cobolt (Stockholm, Sweden) Calypso 491 nm and Jive 561 nm lasers were used with a laser launch built by Solamere Technology Group (Salt Lake City, UT). Images were acquired with Molecular Devices (Sunnyvale, CA) MetaMorph software. The pixel calibration was 165 nm/pixel or 103 nm/pixel, controlled by a tube lens that provided 1x or 1.6x additional magnification. Image processing and analysis was performed using ImageJ.

## Acknowledgments

We thank Dr. Carlos Bustamante and Dr. Daniel Fletcher for their help and generous technical support for C.D.C’s internship and V.I.R.’s graduate research at the University of California, Berkeley. Also, we thank Dr. Daniel Schlieper and Dr. Jan Löwe for the BtubA/B plasmid.

## Author Contributions

The author(s) have made the following declarations about their contributions: Conceived and designed the experiments: CDC VIR FH SDH OM. Performed the experiments:

CDC VIR FH SDH JKP. Analyzed the data: CDC VIR FH JKP DM RDM RL OM. Contributed reagents/materials/analysis tools: RDM OM. Wrote the paper: CDC VIR FH JKP RL RDM OM.

## References

1. Pilhofer M, Ladinsky MS, McDowall AW, Petroni G, & Jensen GJ (2011) Microtubules in bacteria: Ancient tubulins build a five-protofilament homolog of the eukaryotic cytoskeleton. PLoS Biol 9(12):e1001213.

2. Schlieper, D, Oliva MA, Andreu JM, & Lowe J (2005) Structure of bacterial tubulin BtubA/B: evidence for horizontal gene transfer. Proc Natl Acad Sci U S A 102(26):9170–9175.

3. Sontag CA, Sage H, & Erickson HP (2009) BtubA-BtubB heterodimer is an essential intermediate in protofilament assembly. PLoS One 4(9):e7253.

4. Sontag CA, Staley, JT, & Erickson HP (2005) In vitro assembly and GTP hydrolysis by bacterial tubulins BtubA and BtubB. J Cell Biol 169(2):233–238.

5. Jenkins, C, et al. (2002) Genes for the cytoskeletal protein tubulin in the bacterial genus Prosthecobacter. ProcNatlAcadSci U S A 99(26):17049–17054.

6. Pilhofer M, et al. (2007) Characterization of bacterial operons consisting of two tubulins and a kinesin-like gene by the novel Two-Step Gene Walking method. Nucleic Acids Res 35(20):e135.

7. Martin-Galiano AJ, et al. (2011) Bacterial tubulin distinct loop sequences and primitive assembly properties support its origin from a eukaryotic tubulin ancestor. J Biol Chem 286(22):19789–19803.

8. Pilhofer M, Rosati G, LudwigW, Schleifer, KH, & Petroni G (2007) Coexistence of tubulins and ftsZ in different Prosthecobacter species. Mol Biol Evol 24(7):1439–1442.

9. Findeisen, P, et al. (2014) Six subgroups and extensive recent duplications characterize the evolution of the eukaryotic tubulin protein family. Genome Biol Evol 6(9):2274–2288.

10. Desai A & Mitchison TJ (1997) Microtubule polymerization dynamics. Annu Rev Cell DevBiol 13:83–117.

11. Howard J & Hyman AA (2003) Dynamics and mechanics of the microtubule plus end. Nature 422(6933):753–758.

12. Erickson HP &O'Brien ET (1992) Microtubule dynamic instability and GTP hydrolysis. Annu Rev Biophys Biomol Struct 21:145–166.

13. Haeusser DP & Margolin W (2012) Bacteriophage tubulins: carrying their own cytoskeleton key. CurrBiol 22(16):R639–641.

14. Lowe J & Amos LA (2009) Evolution of cytomotive filaments: the cytoskeleton from prokaryotes to eukaryotes. Int J Biochem Cell Biol 41 (2):323–329.

15. Mitchison T & Kirschner M (1984) Dynamic instability of microtubule growth. Nature 312(5991):237–242.

16. Andreu JM & Oliva MA (2013) Purification and assembly of bacterial tubulin BtubA/B and constructs bearing eukaryotic tubulin sequences. Methods Cell Biol115:269–281.

17. Muhlrad, A, Grintsevich EE, & Reisler E (2011) Polycation induced actin bundles. Biophys Chem155(1):45–51.

18. Tobacman LS & korn ED (1982) The regulation of actin polymerization and the inhibition of monomeric actin ATPase activity by Acanthamoebaprofilin. J BiolChem 257(8):4166–4170.

19. Rivera, CR, Kollman JM, Polka JK, Agard DA, & Mullins RD (2011) Architecture and assembly of a divergent member of the ParM family of bacterial actin-like proteins. J BiolChem 286(16):14282–14290.

20. Nishida E & Sakai H (1983) Kinetic analysis of actin polymerization. J Biochem93(4):1011–1020.

21. Polka, JK, Kollman JM, Agard DA, & Mullins RD (2009) The structure and assembly dynamics of plasmid actin AlfA imply a novel mechanism of DNA segregation. J Bacteriol191 (20):6219–6230.

22. van den Ent F, Moller-Jensen J, Amos LA, Gerdes K, & Lowe J (2002) F-actin-like filaments formed by plasmid segregation protein ParM. EMBO J 21(24):6935- 6943.

23. Garner EC, Campbell CS, & Mullins RD (2004) Dynamic instability in a DNA-segregating prokaryotic actin homolog. Science 306(5698):1021–1025.

24. Gayathri P, et al. (2012) A bipolar spindle of antiparallel ParM filaments drives bacterial plasmid segregation. Science 338(6112):1334–1337.

25. Larsen RA, et al. (2007) Treadmilling of a prokaryotic tubulin-like protein, TubZ, required for plasmid stability in Bacillus thuringiensis. Genes Dev 21 (11):1340–1352.

26. Loose M & Mitchison TJ (2014) The bacterial cell division proteins FtsA and FtsZ self-organize into dynamic cytoskeletal Nat Cell Biol 16(1):38–46.

27. Erb ML, et al. (2014) A bacteriophage tubulin harnesses dynamic instability to center DNA in infected cells. eLife 3:3.

28. Takeda M, et al. (2008) Prosthecobacterfluviatilis sp. nov., which lacks the bacterial tubulin btubA and btubB genes. Int J SystEvolMicrobiol 58( Pt7):1561–1565.

29. Cayley S, Lewis BA, Guttman HJ, & Record MT, Jr. (1991) Characterization of the cytoplasm of Escherichia coli K-12 as a function of external osmolarity. ications for protein-DNA interactions in vivo. J MolBiol 222(2):281–300.

30. Alatossava T, Jutt Kuhn A, & Kellenberger E (1985) Manipulation of intracellular magnesium content in polymyxin B nonapeptide-sensitized Escherichia coli by ionophore A23187. J Bacteriol 162(1):413–419.

31. Padan E, Zilberstein D, &Rottenberg H (1976) The proton electrochemical gradient in Escherichia colils. Eur J Biochem 63(2):533–541.

32. Kraemer JA, et al. (2012) A phage tubulinA phage tubulin tubulin assembles dynamic filaments by an atypical mechanism to center viral DNA within the host cell. Cell149(7):1488–1499.

33. Geladopoulos TP, Sotiroudis TG, & Evangelopoulos AE (1991) A malachite green colorimetric assay for protein phosphatase activity. Anal Biochem2(1):112–116.

34. Brouhard GJ, et al. (2008) XMAP215 is a processive microtubule polymerase. cell 132(1):79–88.

